# Reproductive Collapse of Golden Eagles in Oman’s Hyper-Arid Desert: Conservation Implications for Marginal Populations in Extreme Environments

**DOI:** 10.64898/2026.02.26.708261

**Authors:** Jesús Bautista, Elena Bertos, Stuart Benn, Ali Nasser Mohammed Alrasbi, Naser Mubarak Rashid Al Rahbi, José Rafael Garrido, Miguel Ferrer

## Abstract

Hyper-arid ecosystems operate close to physiological tolerance limits, such that relatively small increases in temperature may trigger abrupt and non-linear demographic responses once critical thresholds are exceeded. We analysed long-term climatic trends (1980–2026) and reproductive dynamics of the Golden Eagle (*Aquila chrysaetos*) in the hyper-arid central desert of Oman, one of the southernmost and most climatically marginal populations of the species.

Reproductive and occupancy data were derived from repeated surveys conducted at a minimum of 21 confirmed breeding territories (144 survey visits), complemented by an independent long-term observational dataset (1975–2020; 675 records). Mean annual temperature increased by more than 2 °C over the study period, while precipitation remained persistently low (<40 mm yr◻^1^).

Confirmed reproductive activity declined sharply and collapsed to near zero beyond a narrow thermal threshold (∼28.3–28.6 °C), despite intermittent adult presence. Reproductive activity was strongly negatively correlated with temperature, whereas precipitation showed a secondary effect that did not rescue reproduction once thermal limits were exceeded. Independent demographic observations revealed progressive loss of juveniles and immatures and dominance of isolated adults.

Together, these results provide strong evidence for climate-driven functional extinction sensu reproductive failure, with demographic erosion occurring well before adult disappearance, highlighting extinction-debt dynamics in long-lived desert raptors under ongoing climate warming.

This study has implications for climate adaptation policies in arid regions of the Arabian Peninsula.

## 1. Introduction

Arid and hyper-arid ecosystems are among the environments most vulnerable to contemporary climate change because biological processes often operate close to physiological tolerance limits (IPCC, 2023). Under such conditions, gradual climatic trends may produce abrupt and non-linear biological responses once critical thermal or hydric thresholds are crossed. For long-lived species, these responses may generate a temporal disconnect between environmental degradation and population disappearance, a phenomenon widely described as extinction debt (Tilman *et al*., 1994; Kuussaari *et al*., 2009; Riddell *et al*., 2021).

Large raptors inhabiting desert environments are particularly susceptible to these dynamics. Their reproductive success depends on narrow energetic and climatic windows, whereas adult survival may remain relatively high even under chronic environmental stress (Bahat & Mendelssohn, 1996; Cruz-McDonnell, 2015). Consequently, reproduction is often the first demographic process to fail, while adult presence can persist for decades, masking population collapse if monitoring focuses solely on occupancy. Similar threshold-driven collapses may therefore be increasingly common in marginal populations under rapid warming.

The Golden Eagle (*Aquila chrysaetos*) is one of the most widely distributed raptors globally, yet populations inhabiting the Arabian Peninsula and parts of the Sahara– Sahel region occupy climatically marginal environments near the limits of the species’ tolerance (Watson, 2010; Ellis *et al*., 2024). These desert populations have often been regarded as naturally sparse and irregular breeders, leading to limited conservation concern (Clouet & Goar, 2006). However, recent observations suggest a marked decline in reproductive activity, raising the question of whether this reflects stochastic variability at the range margin or a climate-driven functional collapse.

Central Oman hosts one of the most extreme hyper-arid desert regions historically occupied by golden eagles. Successful breeding was documented during the late twentieth century (Gallagher & Brown, 1982), but subsequent surveys reported declining or absent reproduction despite continued adult presence (Green & Harrison, 2014, 2021). Here, we assess whether long-term climatic warming has exceeded the ecological thresholds required for successful reproduction and whether independent demographic and spatial evidence supports a scenario of climate-driven functional extinction.

## 2. Materials and methods

### 2.1 Study area

The study area comprises the inland hyper-arid desert of central Oman, including Al Wusta Governorate and adjacent interior regions of southern Ad Dakhiliyah and eastern Dhofar (Fig. 1). The region is characterised by extremely low and irregular precipitation, high annual temperatures, intense solar radiation, and minimal vegetation cover. Golden Eagle territories within this area represent the southern climatic margin of the species’ distribution in Arabia.

**Figure 1.**
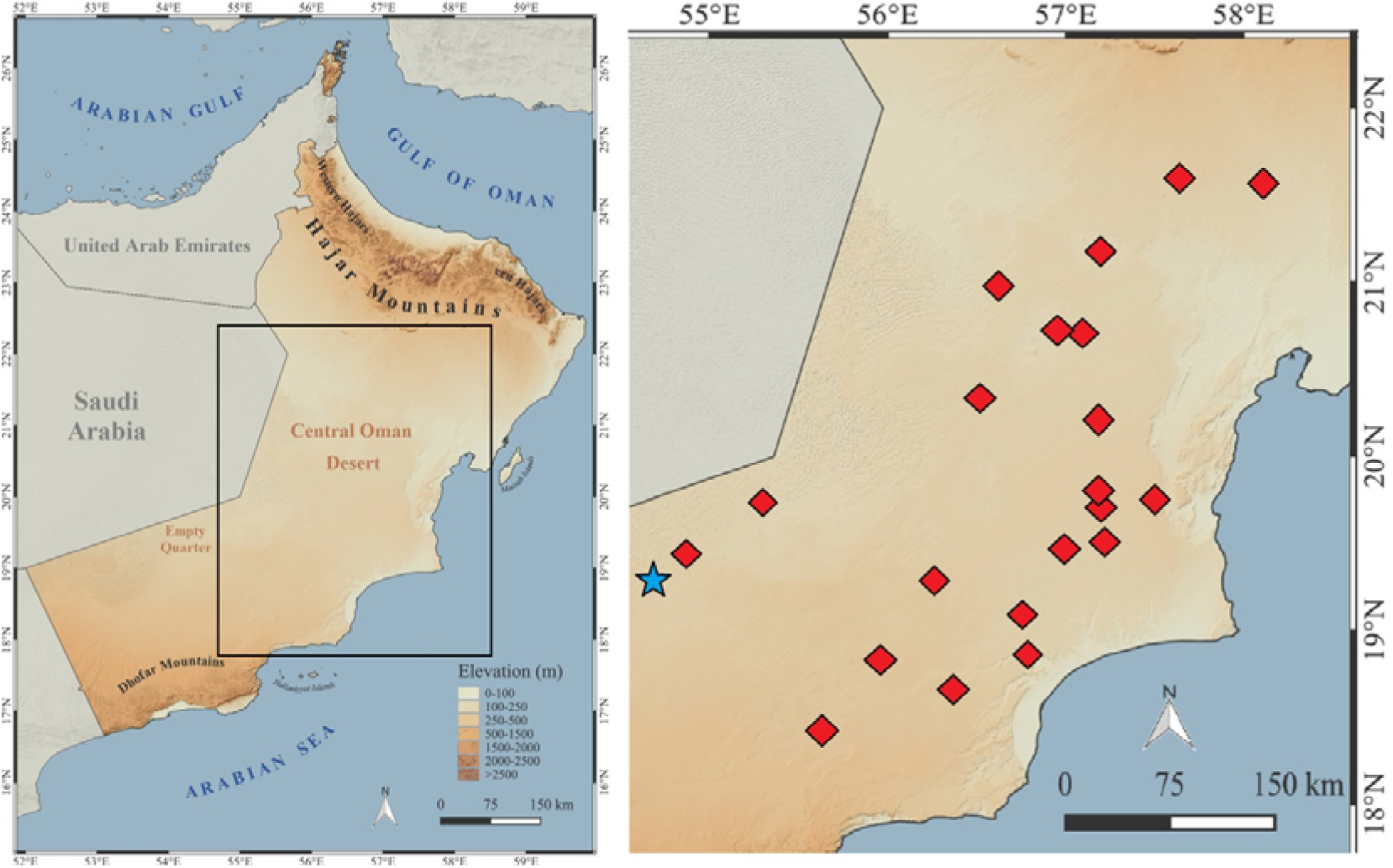
Location of the hyper-arid central desert of Oman showing historical golden eagle breeding territories (red diamonds) analyzed in this study and the position of the Muntasar Oasis (blue star), a key functional habitat used by the species.

### 2.2 Climatic data

Mean annual air temperature (2 m) and total annual precipitation for the period 1980– 2026 were obtained from NASA POWER reanalysis datasets. Climatic variables were extracted at spatial scales representative of golden eagle foraging ranges and were used to characterise long-term directional trends rather than short-term interannual variability (Harmata, 2022; Ellis *et al*., 2024). Potential thermal thresholds were explored through graphical inspection of reproductive output plotted against mean annual temperature.

### 2.3 Reproductive and occupancy data

Reproductive information was compiled from published sources (Gallagher & Brown, 1982; Green & Harrison, 2014, 2021) and from targeted field assessments conducted in 2026. Data were derived from repeated survey visits to known and historically occupied breeding sites within the central desert of Oman.

Across the study period (1980–2026), a total of 144 survey visits were conducted at a minimum of 21 confirmed breeding territories. Reproductive events were conservatively defined as confirmed breeding attempts, including active nests with incubation or chicks observed.

A total of 76 records indicating confirmed reproduction were identified across all data sources. Several of these records referred to the same breeding attempt documented during multiple visits or reported in different sources. To avoid pseudoreplication, these records were consolidated into 70 unique reproductive events, which represent the biological units used in all reproductive analyses.

This conservative approach ensured that each breeding attempt was counted only once, regardless of the number of records documenting it. Occupancy was defined as the detection of at least one adult individual during a survey visit, irrespective of reproductive activity, and was confirmed during 88 visits.

### 2.4 Independent demographic and spatial observational dataset

To complement nest-based reproductive surveys, we analysed an independent long-term observational dataset spanning 1975–2020 and comprising 675 individual records collected opportunistically across the central desert of Oman. Each record included date, location, number of individuals observed, age class (adult, immature, juvenile, chick), and breeding code when applicable.

Within this dataset, records with breeding codes indicating confirmed reproduction (codes ≥11; n = 76) correspond to the same breeding attempts described in Section 2.3. These records were retained in the dataset to characterise long-term demographic structure and spatial use but were not treated as independent reproductive events.

Accordingly, this dataset was analysed separately to assess temporal changes in age structure (presence or absence of juveniles and immatures), demographic erosion, and shifts in the use of key functional habitats (e.g. desert oases), independently of the reproductive output analyses.

### 2.5 Statistical analyses

Relationships between climatic variables (mean annual temperature and total annual precipitation) and biological responses (reproductive events and occupancy rate) were evaluated using Spearman rank correlations, given the small sample size, non-normal distribution of variables, and aggregated nature of the data. Due to strong collinearity between temperature and precipitation, no multivariate models were fitted. Given the rarity of reproductive events, the long lifespan of the species, and the historical nature of the dataset, the objective was not predictive modelling but detection of robust directional and threshold responses consistent across independent datasets. All analyses were conducted in R (R Core Team) using RStudio.

## 3. Results

Reproductive activity (based on unique reproductive events) was concentrated during the 1980s and early 1990s, followed by a progressive decline from the late 1990s onwards. Since approximately 2015, confirmed breeding attempts have been rare or absent, and reproductive activity during the most recent survey period was virtually non-existent.

Reproductive events were strongly negatively correlated with mean annual temperature (Spearman ρ = –0.95, p < 0.001) and strongly positively correlated with total annual precipitation (Spearman ρ = 0.94, p < 0.001; Table 1). Occupancy rate also declined with increasing temperature (Spearman ρ = –0.88, p < 0.01) and increased with precipitation (Spearman ρ = 0.86, p < 0.01). Mean annual temperature and precipitation were themselves extremely strongly and inversely correlated (Spearman ρ = –0.99, p < 0.001).

**Table 1.** Spearman rank correlations between climatic variables (mean annual temperature and total annual precipitation) and biological responses (reproductive events and occupancy rate) of golden eagles (*Aquila chrysaetos*) in central Oman, based on aggregated multi-year temporal bins spanning 1980–2026.

| Variables Compared | Spearman $\rho$ | p-value | Relationship Strength | Ecological Direction |
| --- | --- | --- | --- | --- |
| Temperature vs Reproductive Events | -0.951 | <0.001 | Extremely strong | Higher temperature → fewer reproductive events |
| Precipitation vs Reproductive Events | 0.939 | <0.001 | Extremely strong | Higher precipitation → more reproductive events |
| Temperature vs Occupancy Rate | -0.879 | <0.01 | Very strong | Higher temperature → lower occupancy |
| Precipitation vs Occupancy Rate | 0.855 | <0.01 | Very strong | Higher precipitation → higher occupancy |
| Temperature vs Precipitation | -0.988 | <0.001 | Extremely strong | Inverse climatic relationship |

Graphical analyses revealed a sharp collapse in reproductive output beyond a narrow thermal threshold of approximately 28.3–28.6 °C (Fig. 2a). Beyond this range, reproductive events were rare or absent, whereas adult occupancy persisted at low levels (Fig. 2b), indicating a decoupling between adult presence and reproductive function.

**Figure 2.**
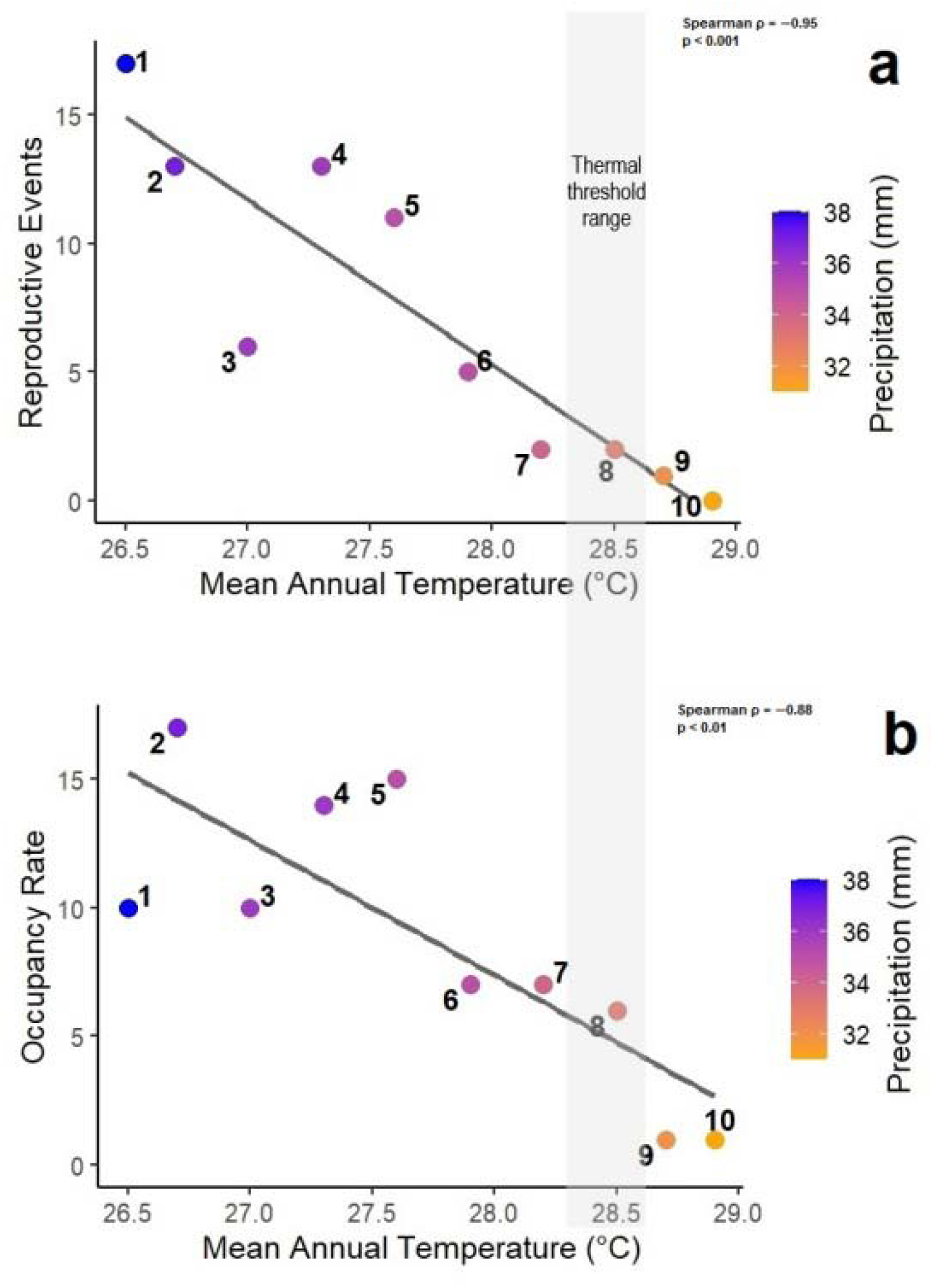
Relationship between mean annual temperature and golden eagle (*Aquila chrysaetos*) reproductive output and occupancy rate in the hyper-arid central desert of Oman. Points represent aggregated multi-year temporal bins spanning 1980–2026. The grey shaded band denotes the approximate thermal threshold range (28.3–28.6 °C).

Analysis of the independent observational dataset revealed marked erosion of population age structure. During the 1980s and early 1990s, records regularly included chicks, juveniles, immatures, and breeding adults, indicating a complete population pyramid. From the late 1990s onwards, juvenile and immature records declined sharply, and after approximately 2005 these age classes were almost entirely absent. Records from 2010–2020 were dominated by isolated adults, indicating a senescent population lacking recruitment (Fig. 3).

**Figure 3.**
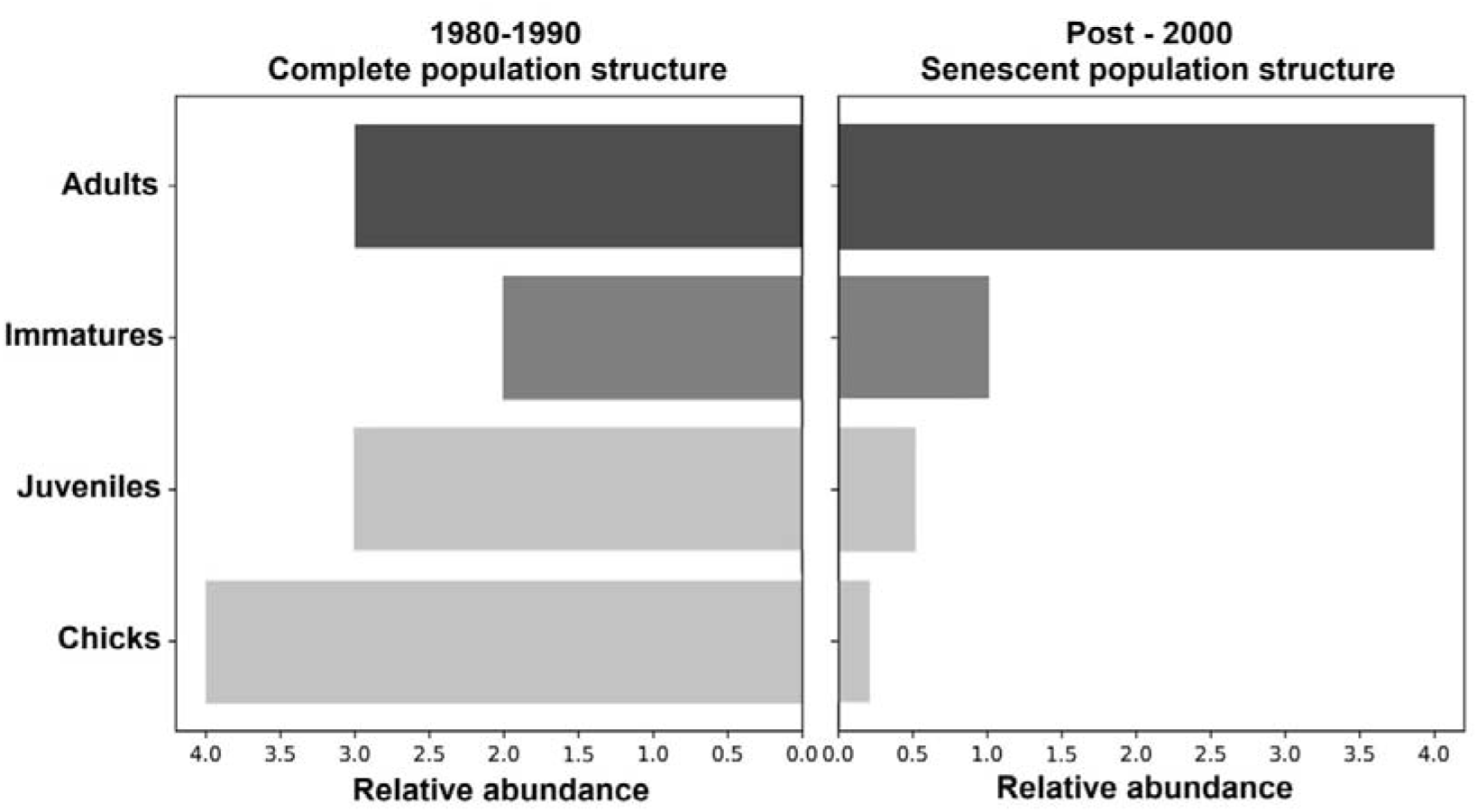
Empirical shift in the age structure of the golden eagle population in central Oman, contrasting a complete population pyramid during 1980–1990 with a senescent, adult-dominated structure after 2000, indicative of long-term recruitment failure and demographic collapse.

Use of the Muntasar oasis declined markedly over time. During the 1980s–2000s, the oasis hosted high numbers of individuals across multiple age classes, acting as a hydric and thermal refuge. After 2010, both visit frequency and numbers of individuals recorded declined sharply, mirroring demographic erosion and indicating loss of landscape functionality.

## 4. Discussion

The decoupling between adult occupancy and reproduction observed in central Oman provides strong evidence for functional reproductive collapse. Despite confirmed adult presence in multiple territories, reproductive output declined sharply and ultimately collapsed, indicating severe and persistent recruitment limitation. Adult Golden Eagles can persist under climatic conditions that no longer permit successful breeding, generating delayed extinction dynamics consistent with extinction debt in long-lived species (Tilman *et al*., 1994; Kuussaari *et al*., 2009).

The existence of a narrow thermal threshold beyond which reproduction collapses suggests a physiological constraint rather than stochastic environmental variability. Experimental and field studies demonstrate that extreme temperatures can compromise gametogenesis, embryo viability, and incubation behaviour in large desert birds (Schou *et al*., 2021; Pattinson *et al*., 2022). In this system, precipitation appears to modulate adult persistence but does not rescue reproduction once critical thermal limits are exceeded. While causality cannot be experimentally demonstrated, the consistency of reproductive, demographic, and spatial evidence across independent datasets strongly supports a climate-driven mechanism. The convergence of these independent lines of evidence greatly reduces the likelihood that the observed reproductive collapse reflects sampling artefacts or short-term environmental stochasticity.

Independent demographic observations further indicate that the population crossed a critical demographic threshold consistent with irreversible decline approximately a decade before adult disappearance. The progressive loss of juveniles and immatures, coupled with the dominance of isolated adults in recent records, reveals a senescent population structure lacking recruitment. Such demographic erosion is characteristic of extinction-debt dynamics, in which long-lived individuals mask population collapse until reproduction has failed for multiple generations.

The concurrent collapse in the use of key desert oases provides additional evidence of functional landscape degradation. Historically, oases such as Muntasar acted as hydric and thermal refuges, supporting multiple age classes and facilitating reproduction in an otherwise extreme environment. The sharp decline in oasis use observed after 2010 suggests that these refuges no longer buffer climatic extremes effectively, further constraining reproductive viability. Loss of access to such functional habitats likely amplifies the effects of rising temperatures, accelerating demographic collapse even before adult disappearance becomes apparent.

Given projected continued warming across the Arabian Peninsula (IPCC, 2023; Malik *et al*., 2024), similar threshold-driven collapses may emerge in other marginal desert populations of large raptors. Populations occupying the warmest and driest portions of species’ ranges are likely to experience functional extinction well before local disappearance is detected through occupancy-based monitoring. These findings underscore the need for conservation assessments that explicitly incorporate reproductive performance, age structure, and extinction-debt dynamics, rather than relying solely on adult presence as an indicator of population persistence.

## 5. Conclusions

Climate warming has driven a functional reproductive and demographic collapse of Golden Eagles in the hyper-arid central desert of Oman. Persistent adult presence masks underlying demographic failure, but the disappearance of juveniles, immatures, and functional refuges indicates a high likelihood of local extinction. These findings highlight the vulnerability of marginal desert populations and the need for conservation assessments that explicitly incorporate reproductive viability and extinction-debt dynamics under ongoing climate change.

## Acknowledgements

We thank all contributors who shared historical data from previous studies in Oman through reports, publications, and personal communications, with particular gratitude to Jens and Hanne Eriksen for species records spanning 1975–2020, complemented by

M.D. Gallagher, M.R. Brown, Mick Green, Ian Harrison, and Richard Porter. We are grateful to Hamid Saif Hamid Al-Ameri and Suleiman Saeed Suhail Al-Harsoussi for their key contributions to the 2026 field study, and to Saleh Mohammed Saleh Alkaabi, Maia Sarrouf Willson, and Bashaar Zaitoon for assistance with fieldwork, logistics, and permitting. We acknowledge the Environment Authority of Oman for providing permits and institutional support, and the IUCN Regional Office for West Asia (ROWA) and the IUCN Centre for Mediterranean Cooperation (IUCN Med) for institutional backing.

## Data availability

Data supporting the findings of this study are available from the corresponding author upon reasonable request, subject to conservation, ethical, and permitting constraints from the Environment Authority of the Sultanate of Oman.

## References

Bahat, O., Mendelssohn, H., 1996. The long-term effect of precipitation on the breeding success of golden eagles Aquila chrysaetos in desert environments. In: Meyburg, B.-U., Chancellor, R.D. (Eds.), Eagle Studies. World Working Group on Birds of Prey, Berlin, pp. 195–202.

Clouet, M., Goar, J.-L., 2006. L’aigle royal Aquila chrysaetos au sud du Sahara. Alauda 74, 441–446.

Cruz◻McDonnell, K.K., 2015. Negative effects of rapid warming and drought on reproductive dynamics and population size of an avian predator in the arid southwest. PhD thesis, University of New Mexico, UNM Digital Repository, Albuquerque, NM, USA.

Ellis, D.H., Bautista, J., Ellis, C. (Eds.), 2024. The golden eagle around the world. Hancock House Publishers, Surrey, British Columbia, Canada.

Gallagher, M.D., Brown, M.R., 1982. The golden eagle breeding in Oman, eastern Arabia. Sandgrouse 4, 100–106.

Green, M., Harrison, I., 2014. Surveys of breeding golden eagles Aquila chrysaetos in the Sultanate of Oman, 2010 and 2014. Confidential report for Ministry of Environment and Climate Affairs, Muscat, Oman.

Green, M., Harrison, I., 2021. The status of golden eagles in Oman. Sandgrouse 43, 1–17.

Harmata, A.R., 2022. Home range and use area sizes of territorial golden eagles tracked with tail◻mounted satellite transmitters. Journal of Raptor Research 56, 479–483.

IPCC, 2023. Climate Change 2023: Synthesis Report. Intergovernmental Panel on Climate Change, Geneva, Switzerland.

Kuussaari, M., Bommarco, R., Heikkinen, R.K., et al., 2009. Extinction debt: a challenge for biodiversity conservation. Trends in Ecology & Evolution 24, 564–571.

Malik, A., Stenchikov, G., Lelieveld, J., et al., 2024. Accelerated historical and future warming in the Middle East and North Africa. Journal of Geophysical Research: Atmospheres 129, e2024JD041625.

Pattinson, N.B., van de Ven, T.M.F.N., McKechnie, A.E., & Cunningham, S.J., 2022. Collapse of breeding success in desert◻dwelling birds within a single decade. Frontiers in Ecology and Evolution 10, 842264.

Riddell, E.A., et al., 2021. Exposure to climate change drives stability or collapse of desert vertebrate populations. Science 371, 633–638.

Schou, M.F., Bonato, M., Engelbrecht, A., et al., 2021. Extreme temperatures compromise male and female fertility in a large desert bird. Nature Communications 12, 666.

Tilman, D., May, R.M., Lehman, C.L., & Nowak, M.A., 1994. Habitat destruction and the extinction debt. Nature 371, 65–66.

Watson, J., 2010. The golden eagle. 2nd ed. T & AD Poyser, London, UK.

